# Effects of Soft Encapsulation on the Receive Performance of PMUTs for Implantable Devices

**DOI:** 10.1101/2025.04.07.646993

**Authors:** Andrada I. Velea, Raphael Panskus, Benedikt Szabo, Vera A. -L. Oppelt, Lukas Holzapfel, Cyril B. Karuthedath, Abhilash T. Sebastian, Thomas Stieglitz, Alessandro S. Savoia, Vasiliki Giagka

## Abstract

Recent studies present ultrasound (US) as a promising candidate for powering implantable devices, requiring integrated and encapsulated receivers to ensure longevity. Conventional hermetic packaging can hinder acoustic transmission, making polymer-based approaches desirable. This study evaluates how polymers commonly used for implants (i.e., thermoplastic polyurethane, parylene-C, medical-grade silicones, and polyimide) affect the receive performance of piezoelectric micromachined ultrasound transducers (PMUTs). Simulations and measurements between 1 and 7 MHz show transmission coefficients above 94 % for material thicknesses in the nm and μm ranges. A theoretical analysis of the mechanical properties guides material selection for later PMUT encapsulation, focusing on polyurethane, parylene-C, and two medical-grade silicones (MED-1000, MED2-4213). In a complete system comprising encapsulated PMUTs, mechanical and acoustic properties, along with interface mismatch between the encapsulation and the PMUTs, influence the receive performance of the devices. Finite element modeling (FEM) and measurements evaluate the impedance and receive sensitivity of encapsulated PMUTs. The results show that residual stress or higher stiffness in some polymers, reduces the receive sensitivity, an effect not evident from only analysing the acoustic transmission through coatings. However, this study demonstrates that upon careful consideration of the acoustic and mechanical properties as well as thickness selection, polymers commonly used for implantable devices can effectively be used for PMUT encapsulation.

## I. Introduction

Advances in the field of implantable devices focus on miniaturization, wireless powering, efficient energy transfer, and highly specific neuromodulation. For wirelessly powered implants, ultrasound (US) has emerged as a promising candidate, particularly for deep-seated implants, due to its efficient propagation through biological tissue. Research shows that US is able to communicate with [1] and deliver high power levels to millimeter-sized (mm-sized) implants at depths over 10 cm inside the human body [2], [3], making it a promising alternative to radio frequency (RF) and inductive coupling methods [4]. Another important advantage is the high (720 mW cm^−2^) FDA-approved limit for the spatial-peak temporal average acoustic intensity (*I*_*SP T A*_) [5]. Using US for powering requires at least two transducers: a transmitter (TX) and an implant-integrated receiver (RX). To enable further miniaturization, micromachined US transducers (MUTs) are currently being investigated. For implantable devices, packaging is crucial, and conventionally, hermetic (either metal or ceramic) cases are being used [6]. However, US transducers require an acoustically conductive medium for optimal sound wave propagation. Metal or ceramic cases (filled with dry gas) hinder signal transmission due to their higher characteristic acoustic impedance, compared to water or biological tissue, causing strong reflections. In an attempt to overcome this, ceramic packages, filled with polydimethylsiloxane (PDMS), and with thin metal lids have been proposed for PZTs to enable ultrasonic coupling while preserving hermeticity [7]. However, the assembly process is cumbersome, and both ceramics, and thin metal layers are prone to cracking, leading to failure and a decreased lifetime of devices. More recent studies [8] and [9] have explored the use of conformal coatings for implantable devices, demonstrating stablity even after in-vivo implantation. Within this frame, investigating similar polymers for the encapsulation of transducers is, therefore, essential. For MUTs in particular, not only do the bulk properties of the encapsulation materials directly influence longitudinal wave propagation, but shear properties and viscoelasticity also play a critical role on the transducer flexural vibration [10]. Hybrid hermetic-soft approaches were proposed for CMUTs in implant applications, where a soft material couples the transducer to the hermetic package [11]. However, this method can still cause unwanted multiple reflections between the soft and hard layers, leading to additional loss, and thus, a polymer-based encapsulation is a more promising approach for ultrasonically powered implants. However, variations in material properties, thicknesses, and deposition methods, can affect the received signal on the RX transducer. This paper systematically analyzes how different polymers affect the receive performance of MUTs by considering the acoustic properties of the materials, their mechanical influence and acoustic mismatches between the soft polymer layers and the harder MUT layers.

The paper outlines the PMUT structure, encapsulation processes, and simulation/ experimental methods in II. Methods. It compares materials and devices with simulations, presents the measurement setup and sample characterisation results in (III. Results and Discussion), and concludes the study in IV. Conclusion.

## II. Methods

### A. Piezoelectric Micromachined Ultrasonic Transducers (PMUTs)

2×2 cm^2^ arrays of piezoelectric-based MUTs (PMUTs) built on Cavity Silicon on Insulator (C-SOI) wafers (Okmetic, Vantaa, Finland), were used. Each device comprises 20 736 cells, each with a diameter of 70 μm and spaced at a 120 μm pitch. The cells are grouped in 16 independently addressable elements (5×5 mm^2^), labeled from A to P, each comprising 1296 cells (**Fig. 1(a)**). The basic structure of a PMUT cell, shown in **Fig. 1(a)**, consists of a C-SOI substrate with a silicon dioxide (*SiO*_2_) passivation layer. The vibrating element (i.e., aluminium nitride (*AlN*) layer) is sandwiched between a bottom molybdenum (*Mo*) and a top aluminium (*Al*) electrode. Finally, a silicon nitride (*Si*_3_*N*_4_) layer acts as passivation for the entire array [3]. The PMUTs exhibit a resonance frequency between 4.5 MHz and 4.7 MHz in air, and 2.9 MHz in water.

**Fig. 1.**
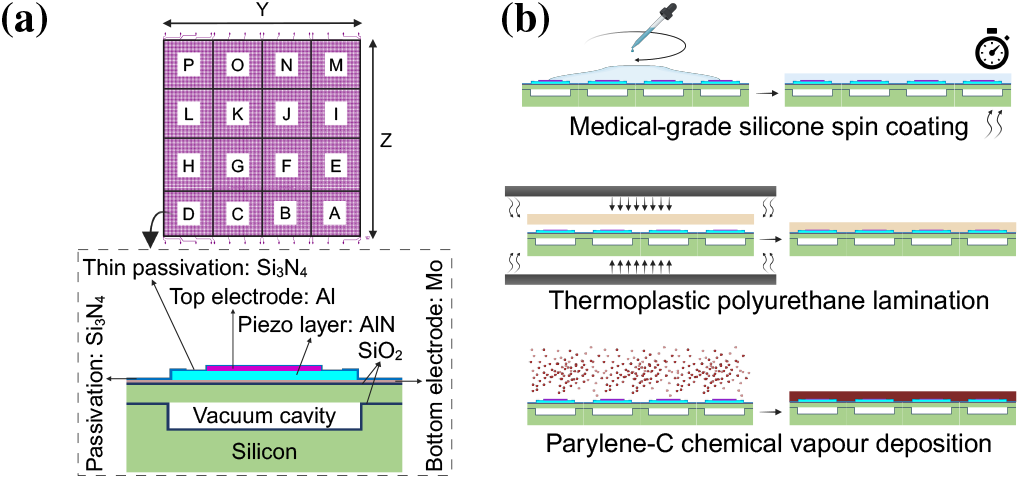
Test structures and encapsulation methods (*Created with BioRender*.*com*). **(a)** Top-view of the PMUT array layout including distribution of the 16 elements and a cross-section schematic representation of one PMUT cell. **(b)** Description of the encapsulation processes used (i.e., spin coating for medical-grade silicones, lamination for thermoplastic polyurethane, and chemical vapour deposition for parylene-C).

### B. Encapsulation materials and processes

For the material investigations, different classes of polymers were considered: thermoplastic polyurethane (TPU) (Platilon 4201 AU, Covestro AG, Germany), medical-grade thermoplastic polycarbonate polyurethane (PCU) (Bionate II 80A, from DSM, The Netherlands), medical-grade silicones (two-component: MED2-4213 and one-component: MED-1000, NuSil Technology LLC, USA), parylene (type C, Galentis S.r.l., Italy) as well as polyimide (Fujifilm LTC9320, Belgium). Upon preliminary theoretical and experimental analysis of the individual materials, for the encapsulation of PMUTs, the material choice has been limited to Platilon 4201 AU TPU, Parylene-C, MED-1000, and MED2-4213 silicones. The encapsulation procedure for TPU, consisted in laminating 25 μm thick material sheets using a cleanroom-compatible thermocompression process at 6–7 bar and 160 ^*°*^C. Due to the thermoplastic nature of the material and the low topography of the PMUTs, the TPU lamination process has the potential of providing reliable tens of μm-thick conformal coatings. The used temperature and pressure does not affect the structural integrity of PMUTs [12]. For parylene-C, 5 μm were deposited by means of chemical vapour deposition (CVD), where parylene monomers were deposited onto the surface of interest at 30 ^*°*^C and 25 μbar. The CVD process offers high conformal coatings even when a few μm-thick layers are desired. Silicones were spin-coated to a target thickness of 45 μm and 37 μm, for MED-1000 and MED2-4213, respectively. Even if the topography of the PMUTs is not significant, a spin-coating process may still lead to non-uniformity or potential defects (presence of gaps between the PMUT and the coating) of the layers.

### C. Acoustic characterisation of coatings

We analysed the acoustic properties of the selected coating films by simulating and measuring the acoustic wave transmission coefficient (T) through polymers with thicknesses ranging between 100 nm and 5 mm. Numerical simulations based on [13], and described in detail in [14] assume continuous transmission of planar waves, with pressure and particle velocity continuity at each boundary. As illustrated in **Fig. 2(a)**, the model estimates T through various material films, in a semi-infinite water medium, taking into account reflections and attenuation of the acoustic wave. The material properties used in the simulations are listed in **Table I**. Density, *ρ*, speed of sound, *c*, acoustic impedance *Z*, and bulk attenuation, *α* at 5 MHz were measured according to the methodology described in [15] using 405×405×2 mm^3^ material slabs. The thickness was chosen such that the material samples were significantly thicker than the wavelength, and thus the parameters of interest were accurately extracted. Since for parylene-C and polyimide, such thicknesses were unachievable, the values were based on previous literature. For the measurements, material films were placed inside a water tank, between an unfocused 5 MHz transmit (TX) transducer (V309-SU, Olympus, Tokyo Japan) and a 1 mm needle hydrophone (NH1000, Precision Acoustics, Dorchester, UK) (**Fig. 2(b)**). The TX transducer was driven using sinusoidal pulses, modulated by a Hann window, generated by a function generator (Keysight 33600A, California, USA). The pulse width was 20 μs with 4 μs rising and falling edges, a pulse repetition frequency (PRF) of 20 Hz and the peak-to-peak amplitude of 1 V. The signal amplified using an RF amplifier (350L, E&I, New York, USA) with a 47 dB nominal gain. First, reference measurements (without a polymer film) were taken, for frequencies between 1 MHz and 7 MHz, in steps of 1 MHz which include the resonant frequencies of the PMUTs, both in air and in water, at a distance of 10 cm. To record the acoustic pressure through materials and convert it into an electrical signal, the needle hydrophone was connected through a DC coupler (Precision Acoustics, Dorchester, UK) to an oscilloscope (RTA4004, Rhode&Schwarz, Munich, Germany). On the walls of the water tank, particularly behind the hydrophone, under the sample, and lateral to the transducer, sample, and hydrophone, absorbers (VK-76000, Gampt, Merseburg, Germany) were placed in order to reduce reflections from the tank walls. To further minimize errors and maintain consistent measurement conditions, the samples were positioned using a motorised 3D stage (SFS630, Gampt, Merseburg, Germany) at an equal 10 cm distance both from the transducer and the hydrophone. This minimizes reflections, preventing interference with the measured signal. To verify this, a 7 MHz single-cycle pulse was sent through the polymer and the time delay between the first reflection and the main recorded signal was monitored. The hydrophone-transducer distance was adjusted to ensure a time delay over 20 μs, which is the pulse width used during the experiments. Pressure through the polymer samples was then measured and the transmission coefficient was calculated by subtracting the material measurements from the reference measurements (assumed to have a normalized transmission coefficient of 1), at each frequency.

**TABLE I.**
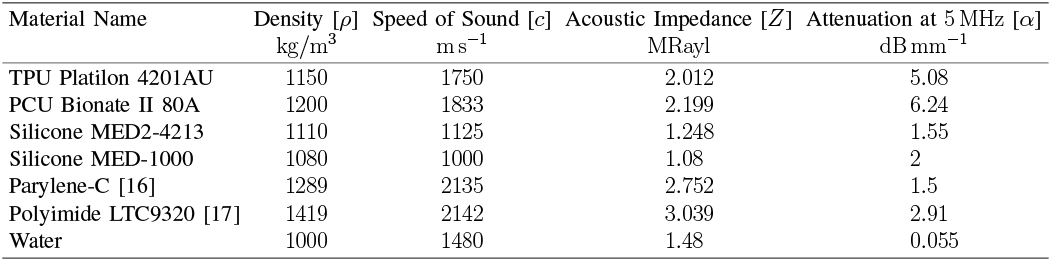
Material properties used for simulations.

**Fig. 2.**
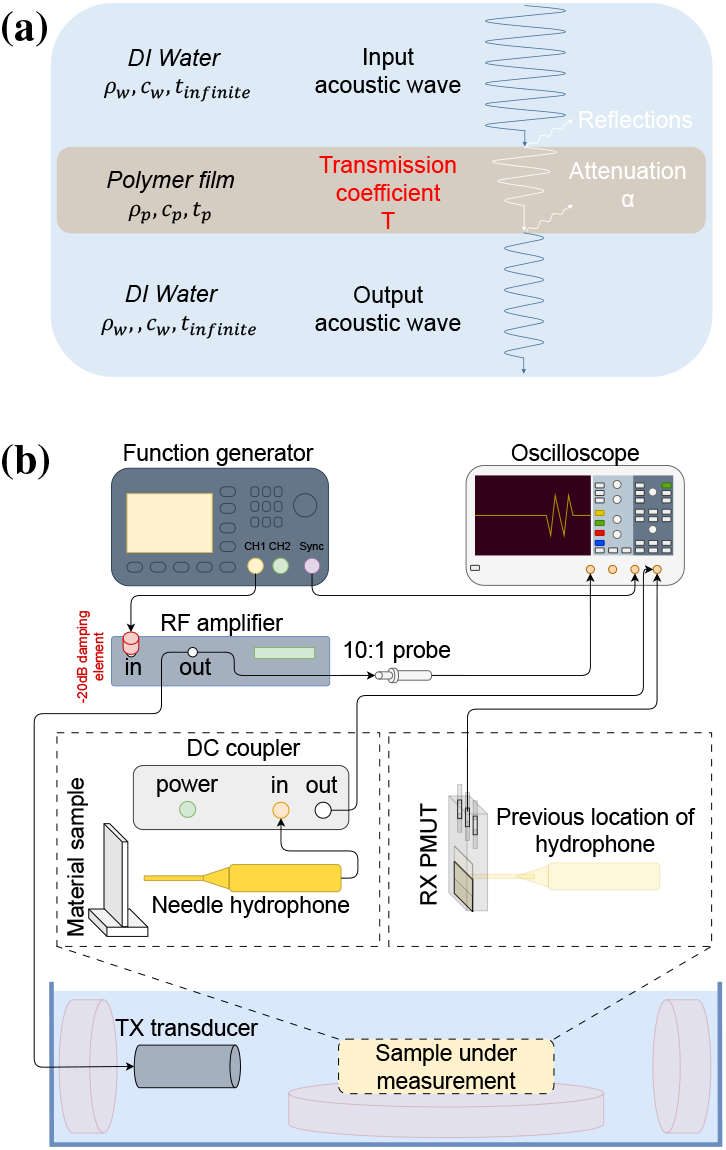
Simulation model and test setup used for evaluating the transmission coefficient. **(a)** Simulation model for the polymer film materials, where *ρ* represents the density, *c*, the speed of sound, *t*, the thickness, *α*, the attenuation, and *T* the simulated and measured transmission coefficient. **(b)** Experimental setup used for measuring the transmission coefficient through polymers as well as the open circuit voltage (OCV) of coated and uncoated PMUTs (*Created with draw*.*io*).

### D. Acoustic characterisation of encapsulated PMUTs

#### 1) Finite Element Modelling (FEM)

Finite element modeling (FEM) using ANSYS (Ansys Inc., Canonsburg, PA, USA) was performed to predict the behavior of coated and uncoated PMUTs in terms of electrical impedance and receive sensitivity. A 3D FEM model of a periodic layout of circular PMUT cells arranged in a square grid pattern, with the layer stack described in II-A, was implemented using ANSYS APDL. The top of the cells is encapsulated, while the 660 μm-thick bulk Si of the PMUT is in contact with a 1.55 mm-thick FR4 printed circuit board (PCB). Both the PMUT cells and the FR4 layer are coupled to a fluid medium (water). The solid and fluid materials were meshed using hexahedral SOLID185 and FLUID30 elements, respectively, while SOLID226 elements were used to model the piezoelectric transduction effect. To approximate coupling to an infinite propagation medium, total reflection was assumed at the boundaries of the top and bottom fluid domains. The acoustic attenuation in the encapsulation layer was modeled using a linear viscoelastic model, employing a Prony series representation of bulk relaxation behavior. The coefficients of the Prony series were fitted to match the experimentally estimated bulk attenuation around 5 MHz, following established methods for frequencydependent damping in soft polymers [10][18]. To calculate the electrical impedance, a voltage generator modeled using the CIRCU94 element was applied to the top and bottom electrodes of the PMUT cells. Harmonic analyses were performed with a uniform voltage excitation over the 0.5–7 MHz range for the bare PMUT in air and water, as well as for different encapsulation materials under water-coupled conditions. To calculate the receive response, a uniform pressure excitation was applied at the center of the fluid domain to generate a plane wave directed towards the PMUT. Harmonic analyses were then conducted over the same frequency range for each encapsulation material in water-coupled conditions. Platilon TPU (25 μm), parylene-C (5 μm), MED-1000 (45 μm) and MED2-4213 (37 μm) silicones were considered for FEM modelling, as these were later used for PMUT encapsulation and analysis. The acoustic and mechanical properties used in the simulations have been presented in tables I and II. Additionally, preliminary COMSOL modelling has been employed to simulate the forces at the interfaces between MED-1000 silicone and the PMUT cells. The model uses a 7.43 μm size triangular mesh, and it assumes PMUT cells with the layer stack described in II-A and a 100 μm encapsulation layer on top.

#### 2) Impedance measurements

Impedance on uncoated and coated PMUTs was measured with 1601 points per measurement, between 1 MHz and 7 MHz using an impedance analyzer (Keysight E4990A, California, USA). A total of 144 individual partitions from nine uncoated and wire bonded samples were measured in air and deionized (DI) water (an excellent coupling medium with acoustic impedance similar to soft tissue). For the water-coupled measurements, each PMUT sample was placed inside the water tank, facing an absorber to prevent reflections from influencing the measurements. Next, the samples were coated with different polymers, as described in II-B, and remeasured. Each polymer was deposited on a separate uncoated sample (a total of 16 partitions per polymer were measured). Encapsulation with Platilon TPU was not possible on wire-bonded PMUTs due to potential damage of the wires during the encapsulation process. Thus, the sample was first coated, then wire bonded and its impedance measurements were compared to other uncoated devices. Since PMUTs show broadband behavior in DI water, air-coupled impedance measurements were primarily used for comparison. An initial system calibration was required, considering additional components, such as cables and PCB, thus increasing the signal-to-noise ratio. For the uncoated devices, measurements were first averaged across partitions within each device. The final plotted result represents the average of these partition-averaged values across all uncoated samples, along with the standard deviation. For the coated devices, since each coating corresponds to a single device, measurements were averaged across partitions, and this average was plotted with its associated standard deviation.

#### 3) Receive sensitivity measurements of PMUTs

Measurements on uncoated and coated PMUTs were conducted to evaluate the effect of soft encapsulation on the received energy. The setup in **Fig. 2(b)** is the same as for the material characterization, but the material samples were replaced by functional PMUTs. First, the output pressure of the TX transducer (described in II-C) was evaluated in its far field, at a distance of 25 cm (where the PMUT was later placed for evaluation), for frequencies between 1 MHz and 7 MHz, in steps of 0.2 MHz. Manual alignment was employed to find the maximum output pressure of the transmitter. The input impedance of the oscilloscope was set to 1 MΩ. Additionally, a 2D scan measurement was performed to further evaluate any changes in the beam profile of the TX transducer, over the frequencies of interest. Finally, the recorded voltage (for both types of measurements) was converted into pressure, considering the sensitivity of the needle hydrophone used (**Fig. 2(c)**). To measure individual partitions, the PMUT was moved in the Y and Z directions with respect to the TX transducer, using an automated 3D stage. The measured open circuit voltage (OCV) of each PMUT partition was recorded, once, for each frequency. Given the measured OCV and the pressure at the PMUT location, the RX sensitivity of each partition was calculated by dividing the two values, and the average over all partitions, and the respective standard deviation were plotted. Since more uncoated devices were measured, the averaging and standard deviation were calculated and plotted following the same principle described in II-D2. Small variations in the measurement setup, such as water level and sample positioning, could introduce errors. To address these, the setup was further characterized, by evaluating lateral and angular misalignments between the TX transmitter and PMUTs, as well as analysing the measurement errors.

## III. Results and Discussion

### A. Acoustic characterisation of coatings

The selected thickness range for evaluating the transmission coefficient reflects typical values for neural interface substrates and encapsulation. **Fig. 3(a)** highlights key dependencies for the frequency range of interest: material attenuation differences at the same thickness and distinct resonance peaks in thicker films due to changing resonance conditions. For example, for 5 mm compared to 1 mm thicknesses, both peak positions and widths, change. A preliminary classification for the materials can be derived with MED2-4213, which overall has the highest transmission coefficient due to the relatively low attenuation and minimal acoustic impedance mismatch with water (**Table I**). It is followed by parylene-C, MED1000 (although this can alternate for low frequencies and thin layers, it can be assumed that the difference between the two materials is relatively small), polyimide, and polyurethane (Platilon and Bionate). For thicknesses in the nm-range, the transmission coefficient remains above 99.9 %,with negligible losses. Even for μm-range, although T has a tendency to drop for some materials, T is always above 90 % at a maximum frequency of 7 MHz. As shown in **Fig. 3(b)**, the measured and simulated transmission coefficients align. The thicknesses for the material samples were different and dependent on the deposition processes. The larger variations observed for silicones are likely due to the non-uniformity of the spincoated layers. This was most notable for MED2-4213, where white light interferometry showed μm-scale surface roughness. Yet, T remained above 94 % in all cases.

**Fig. 3.**
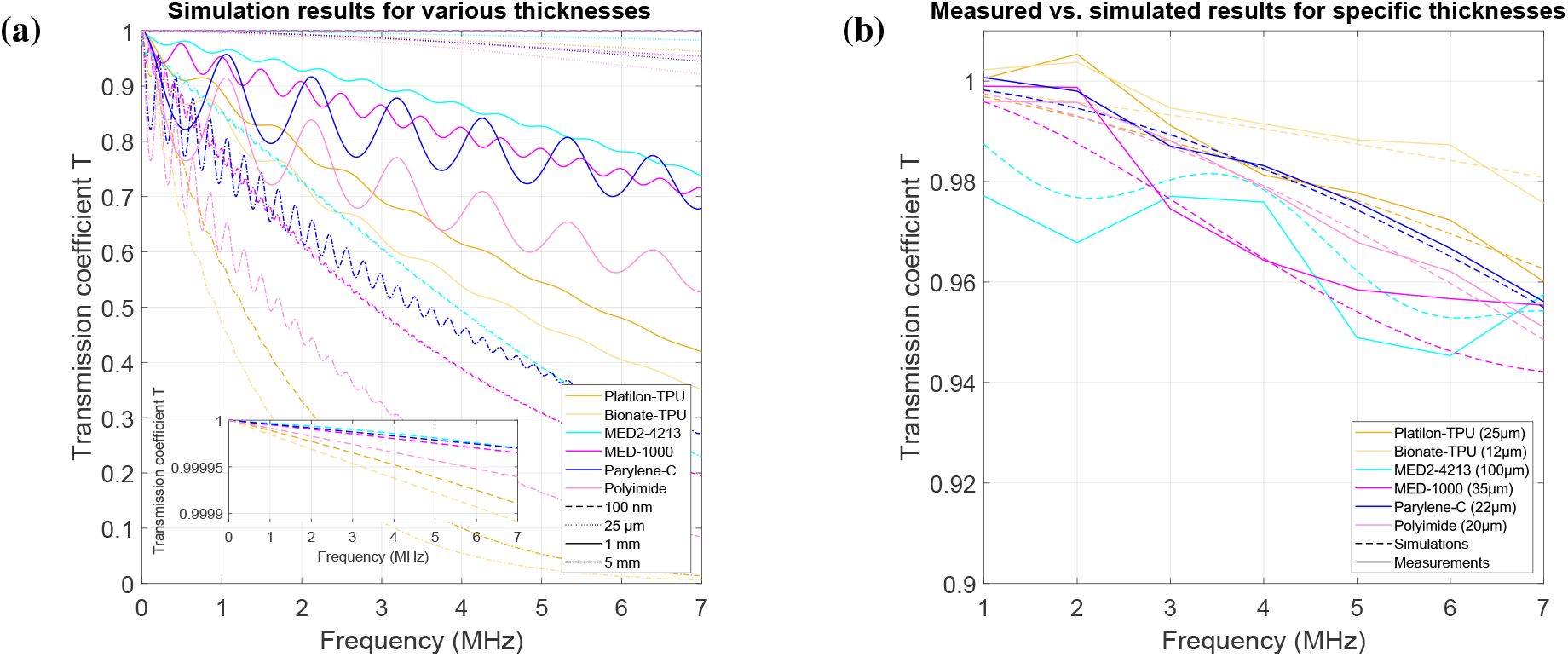
Acoustic characterisation of materials. **(a)** Simulated transmission coefficient through polymers thicknesses from 100 to 5 mm. An inset for thin layers (100 nm) was added. The transmission coefficient decreases with increasing thicknesses and frequencies. The resonance peaks and changes in resonance conditions are visible, particularly for thick layers. **(b)** Measured transmission coefficient juxtaposed against simulations for thicknesses and materials of choice. The transmission coefficient in all considered cases is above 94 %.

### B. Mechanical analysis of coatings

Mechanical properties and residual stress inside polymers may limit the movement of PMUT cells, potentially reducing their performance. The two silicones investigated have different crosslinking mechanisms: MED-1000, a one-component self-curing silicone, cures rapidly in the surrounding humid atmosphere, but exhibits a shrinkage effect, potentially affecting membrane deflection. MED2-4213, a two-component silicone, cures without shrinkage and thus without internal stresses. Preliminary COMSOL modelling shows that the residual stresses of MED-1000 silicone layer are 4 kPa, leading to an upward lift of the membranes of approximately 4 nm. To support the future selection of materials and thicknesses for PMUT encapsulation, the tensile strength, Young’s modulus as well as elongation properties (**Table II**), were analysed, from a theoretical perspective. For example, polyimide is significantly stiffer, with a Young’s modulus 3–4 orders of magnitude higher than silicones and TPU and elongation one order lower than the rest of the materials. Since it is expected to restrict the movement of the PMUT cells, thus significantly reduce the receive sensitivity of the devices, it was excluded from further studies in this paper.

**TABLE II.** List of mechanical properties of chosen polymer families.

| Material Name | Tensile Strength<br>MPa | Young's Modulus<br>MPa | Elongation<br>% |
| --- | --- | --- | --- |
| Polyurethane [19] | $24.8 \pm 1.7$ | $33.4 \pm 3.0$ | Up to 336 |
| Silicone [20] | 6.2 | $0.3 \pm 0.2$ | Up to 600 |
| Parylene-C [21], [22] | 55.16 | 2750 - 4500 | Up to 200 |
| Polyimide [20] | 392 | 8830 | Up to 30 |

### C. Electrical impedance analysis

Air- and water-coupled electrical impedance was evaluated by means of FEM and measurements on coated and uncoated PMUT samples, and the results are shown in **Fig. 4**. The air-coupled simulations in **Fig. 4(a)** indicate a shift in the resonance peaks for the encapsulated PMUTs, from ∼ 4.7 MHz to frequencies between 2.1 MHz and 2.5 MHz for silicone and TPU-coated PMUTs, and ∼ 4.2 MHz for parylene-C-coated PMUTs. This is due to the fact that the additional layers change the resonator characteristics of the PMUT by adding additional mass and stiffness to the membranes. A stiffer coating, such as parylene-C, often indicates a higher resonance frequency, compared to softer polymers, such as silicones and TPU (**Fig. 4(a)**). The dampened peaks can be attributed to the viscoelasticity of the polymers, which in turn can lead to energy dissipation within the coating. The slightly wider resonance peaks (for TPU and parylene-C-coated samples) indicate an improvement in the acoustic matching by providing a better transition between the high PMUT and the low air acoustic impedance. A similar effect occurs when the samples are water-coupled. As shown in the simulated phase (inset in first subplot in **Fig. 4(b)**, the resonance frequency drops significantly to values around 2.9 MHz for all considered cases, due to the added load of the surrounding medium. An even more broadband behaviour of the PMUTs can also be observed, indicating a significant improvement in the impedance matching between the PMUTs and water. Although some variations are present, the measured results (second subplots in **Fig. 4(a)** and **Fig. 4(b)**) show similarities to the simulations. The measured air-coupled resonance frequency in **Fig. 4(a)**, for the 45 μm thick MED-1000 silicone and 25 μm TPU-coated devices, shifts to lower values, as predicted by the simulations. An additional peak can be observed for the TPU-coated samples (zoomed-in inset in second subplot of **Fig. 4(a)**). This can be attributed to one of the shear mode resonances of TPU, which is more visible also in the simulations phase plot (inset in **Fig. 4(b)**). In **Fig. 4(a)**, it can be noted that the resonance peak for the 5 μm parylene-C-coated devices is highly damped and widened, possibly due to internal stresses in the film which modify the overall stiffness of the membrane, introducing additional damping. An interesting discrepancy arises in the 37 μm MED2-4213-coated samples, where the measured air-coupled resonance frequency is higher than in simulations (**Fig. 4(a)**). This suggests possible inconsistencies between the simulated and actual material properties. Particularly, the speed of sound can be assumed to be higher in reality, which would explain a shift towards higher frequencies for the MED2-4213-coated samples. However, a more likely cause can be the presence of gaps between the PMUT and the encapsulation (i.e. air interface), which could lead to an increase in the resonance frequency. It can also be noted that the standard deviation in the phase plot increases with frequency, both for the air- and water-coupled measurements (insets of second subplots in **Fig. 4(a)** and **Fig. 4(b)**). This is most likely a systematic effect due to a non-perfect system calibration prior to the measurements. Although the scale for the plots in these figures is different for a better illustration of the results, the standard deviation trend is similar. During calibration, the additional components comprising the setup are taken into account, except for the on-chip resistance of the PMUT, which is larger for inner elements, due to the longer traces to the pads. This, in turn, adds to the overall electrical impedance. The water-coupled simulations, shown in the first subplot of **Fig. 4(b)** assume the real-case scenario, where the PMUT was stacked on top of a PCB and surrounded by water. The variations observed in the measured phase plot, (second subplot of the same figure), compared to the simulations can be caused by the on-chip resistance of the PMUTs. Although a correction was applied by subtracting the on-chip resistance (assumed to be the real part of the air-coupled impedance at high frequencies) from the water-coupled measurements, the correction factor may still vary more than estimated. The water-coupled simulations reveal periodic resonance peaks observed in the phase plot (inset first subplot of **Fig. 4(b)**), which can be attributed to the PCB material. Moreover, as previously described, the shear mode resonance peak of the material itself is present around 1 MHz for the TPU-coated sample. However, for the frequencies of interest, this should not have a significant impact on the receive performance of the PMUTs.

**Fig. 4.**
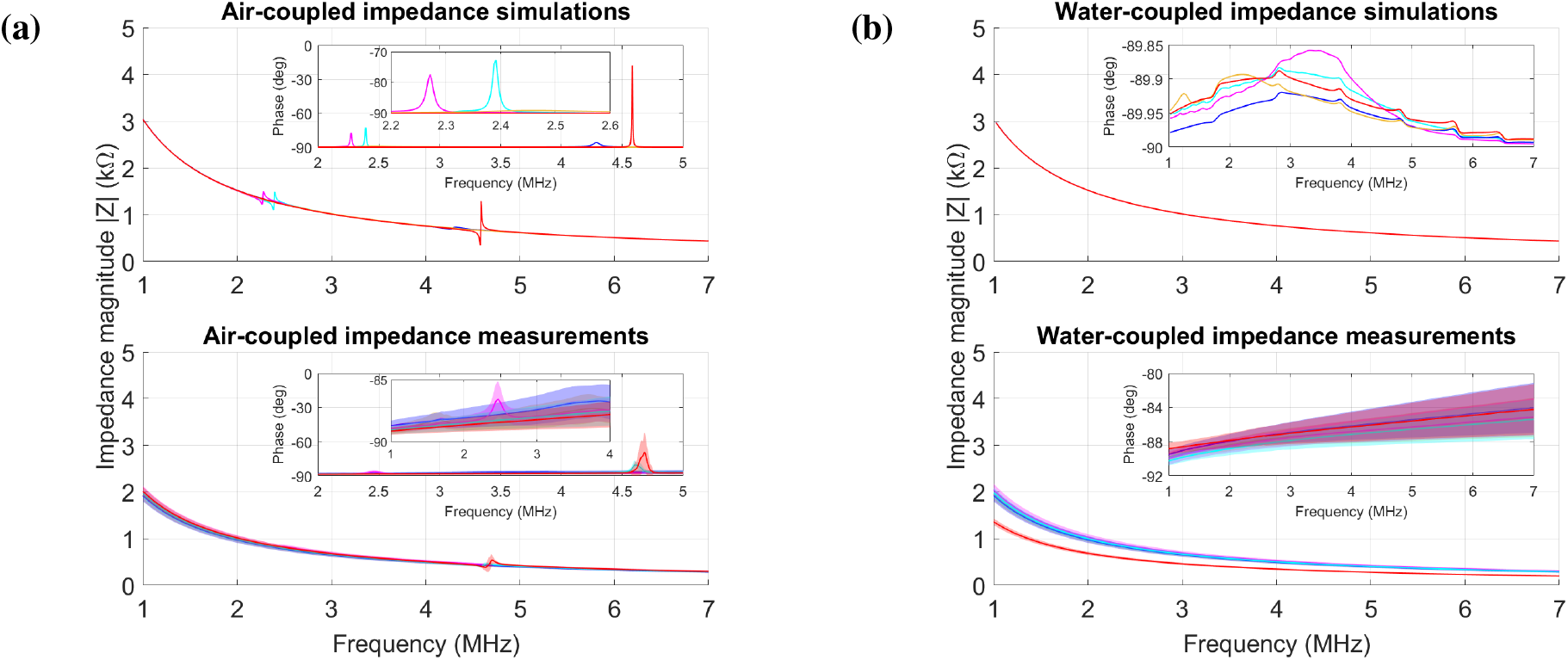
Impedance measurements of coated and uncoated PMUTs. **(a)** FEM simulations of air-coupled electrical impedance. The resonance peaks, together with the frequency shift and dampened peaks for the simulated encapsulated samples can be observed. Air-coupled impedance measurements show higher resonance for MED2-4213-coated samples than simulations, potentially due to the presence of air gaps between the encapsulation and the PMUTs (legend is the same as **Fig. 5(b)** and axes in the second inset are the same as in the first). **(b)** FEM simulations of water-coupled electrical impedance. Due to the broadband behavior of the PMUTs in DI water, the resonance peaks are not clearly visible. The effect of the PCB and the shear mode resonances for TPU-coated samples can be observed. Water-coupled impedance measurements. The large standard deviation in the phase can be attributed to the on-chip resistance of the PMUTs (legend is the same as **Fig. 5(b)**)

### D. Receive sensitivity of uncoated and coated PMUTs

The simulated results, illustrated in **Fig. 5(a)**, show that for all coated PMUT devices, the receive sensitivity is comparable to that of an uncoated PMUT. Small deviations are present, particularly for the parylene-C-coated sample, where a maximum drop in RX sensitivity of ∼ 25 %, at low frequencies can be observed. This drop in the receive sensitivity is also observed in **Fig. 5(b)** and can be attributed to the mechanical properties of the material rather than to the acoustic ones. As shown in **Fig. 3**, the material alone, at the specified thickness can be considered acoustically transparent. As shown in III-C, the thickness mode resonance peaks induced by the PCB material, having the fundamental frequency around 1.5 MHz and all other harmonics at higher frequencies, are also present in **Fig. 5(a)** and **Fig. 5(b)**. Moreover, for the TPU-coated sample, at frequencies around 1 MHz (**Fig. 5(a)**), as also shown in the impedance simulation results in III-C, the shear mode resonance peak can be observed. These however, do not seem to affect the RX sensitivity of the simulated samples. For the measured samples, however, the frequency at which the peaks appear does not accurately match the simulations. This is due to the mismatch between the assumed PCB material properties and the actual values. The slight decrease in the overall recorded sensitivity magnitude (**Fig. 5(b)**) for all samples, compared to the simulations (**Fig. 5(a)**) is expected and assumed to be caused by additional components such as cables, that are not considered in the simulations or even by the error introduced by the measurement setup. If variations occur between samples, this will be reflected in the averaged results. It is important to note that in case some of the partitions under measurement were completely unresponsive due to damage to the membranes or to the wire bonds, they have been excluded from the averaging and, thus, from the final analysis. Compared to the simulations, the trend in the measurement results is generally followed by the PMUTs. For MED2-4213, it can be observed (**Fig. 5(b)**) that the sample exhibits an RX sensitivity similar to the average RX sensitivity of uncoated PMUTs, thus accurately matching the simulations. However, the predicted RX sensitivity for MED-1000-coated PMUTs is not reproduced by the measurement results. The drop in sensitivity as well as large standard deviation (larger than for the rest of the samples) can be attributed to the residual stresses that the material applies to the PMUT cells, as discussed in III-B, thus significantly reducing the measured OCV, which, in turn, results in a decrease in the received pressure. Another important aspect observed in the measured data in **Fig. 5(b)** is the decreased RX sensitivity at particular frequencies. This can be caused by the fact that frequencies above ∼ 4 MHz are outside the frequency bandwidth of the PMUTs, in water. Moreover, frequencies around 1 MHz are also outside the frequency bandwidth of the TX transmitter (an effect shown in **Fig. 5(d)**). Therefore, both the transmit sensitivity of the TX transducer and the receive sensitivity of the PMUT are considerably low. Additional 2D scans of the TX transducer have been carried out to evaluate the changes, not only in the intensity of the transmitted acoustic beam (**Fig. 5(e)**) but also in its profile, evaluated at the Z of maximum measured intensity. As illustrated in **Fig. 5(f)**, the higher the frequency, the narrower the acoustic beam gets. This, in turn, can influence the results obtained while measuring the OCV of the PMUTs. Assuming that misalignment can occur, if the beam, at high frequencies, is narrow or even narrower than the partition under measurement (5 mm by 5 mm), a significant drop in the receive sensitivity can be observed. Additional factors, including measurement error, that can affect the results and cause discrepancies were explored in **Fig. 6**. To quantify the setup error, 10 consecutive measurements were performed on the same sample without changes, and the results indicated a maximum error of ± 3 % (**Fig. 6(a)**). Furthermore, setup changes introduce errors. Measurements on different days, requiring the sample to be removed from the stage or disassembling of the setup between measurements cause OCV fluctuations, even with the same settings (**Fig. 6(b)**). Equally important is the effect of misalignment. In **Fig. 6(c)**, both angular as well as lateral misalignment were evaluated. An angular misalignment, of even a 1^*°*^ could potentially lead to a decrease of up to 20 % in the measured OCV, for the highest measured frequency. For small lateral misalignments the effect does not appear to be as significant. However, it can be noted that a misalignment of 5 mm leads to a decrease in the measured OCV of up to 60 %, for the highest measured frequencies. Similar results have been previously simulated and reported in [23].

**Fig. 5.**
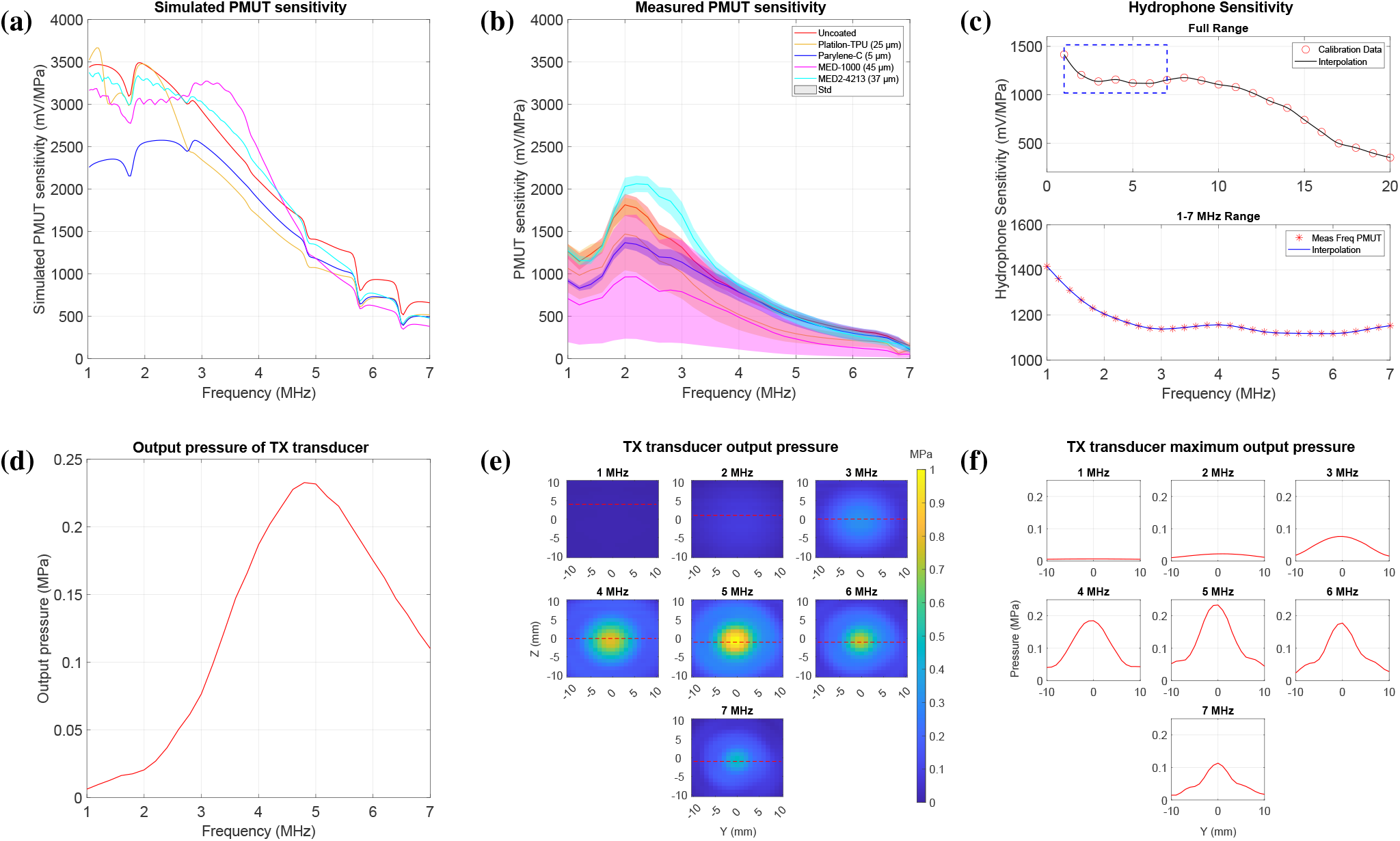
Receive sensitivity of PMUTs. **(a)** FEM simulations (legend is the same as (b)). Coated samples exhibit similar receive sensitivity as uncoated PMUTs. A maximum drop of 25 % in the receive sensitivity at low frequencies can only be observed for a simulated parylene-C-coated sample. **(b)** Receive sensitivity measurements of PMUTs. The measured receive sensitivity of MED-1000 coated PMUTs is significantly lower than predicted in the simulations, potentially due to the residual stress applied by the polymer. **(c)** Sensitivity of a 1mm needle hydrophone, plotted according to the values specified in the datasheet. **(d)** Output pressure of TX transducer measured in its far field, at a distance of 25 cm. The highest output pressure is measured around the resonance frequency of the transducer. **(e)** 2D scan of the output pressure of TX transducer measured in its far field, at a distance of 25 cm. The dotted line represents the area of highest intensity and is then plotted in (f) to illustrate the beam width. **(f)** Width of the TX beam extracted from the 2D scans of the measured output pressure of the TX transducer. The change in output pressure as well as beam profile is visible with the change in frequency.

**Fig. 6.**
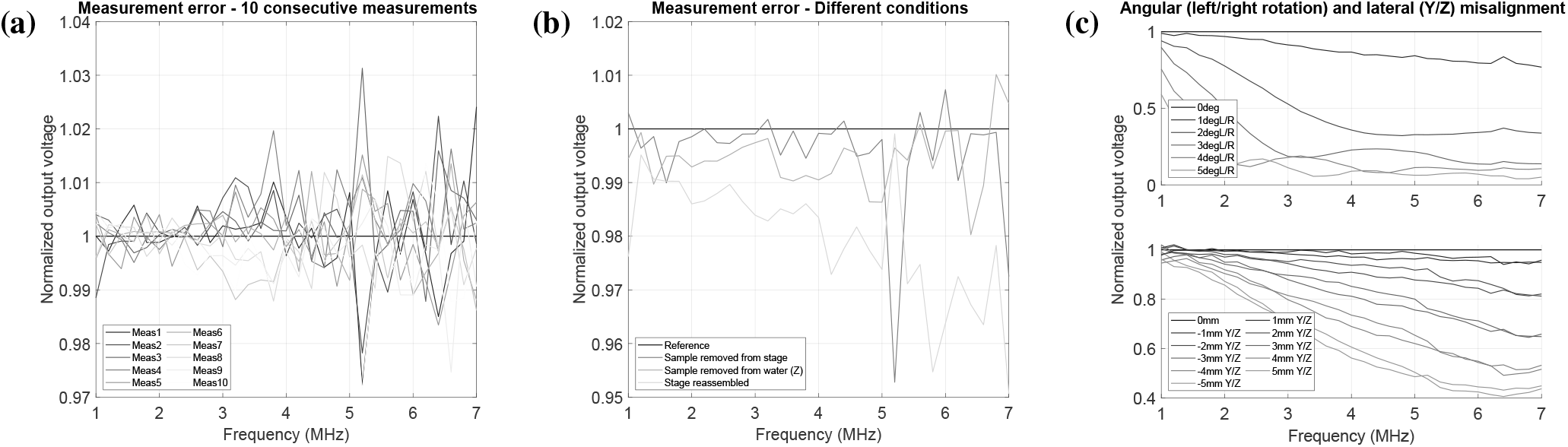
Measurement variability and setup error. (a) Measurement error after 10 consecutive measurements on the same sample. A maximum error of ± 3% was recorded. (b) Normalized single OCV measurements for one uncoated Rx PMUT under different setup conditions. The highest drop is observed when the measurement setup is partly reassembled, between measurements. **(c)** Error from angular misalignments of up to 5^*°*^ in both left (L) and right (R) directions of the Tx transducer relative to the measured PMUT. A decrease up to 20 % in the measured OCV, for 1^*°*^ misalignment, for the highest measured frequency is recorded. Error introduced by lateral misalignments of up to 5 mm and 5 mm both in Y and Z directions. A misalignment of − 5 mm leads to a decrease in the measured OCV up to 60 %, for the highest measured frequencies.

## IV. Conclusion

This study evaluates the receive performance of polymer-coated PMUTs for implantable devices through simulations and measurements. The PMUTs used during the investigations were coated with different types of polymers, commonly used for packaging of flexible implants. From a purely acoustic perspective, for thicknesses in the nm and μm ranges, all materials exhibited a transmission coefficient above 90 %, and above 94 % for selected thicknesses, for frequencies up to 7 MHz. This, in turn, makes them good candidates for applications requiring encapsulated PMUTs. FEM modelling, coupled with extensive measurements on various coated and uncoated samples was employed to estimate the dynamic behaviour of encapsulated PMUTs and evaluate the receive performance of the devices. The results indicated that for the final selection of materials and thicknesses, all polymers exhibit characteristics that could make them suitable candidates for PMUT encapsulation. However, residual stress, particularly for MED-1000 silicones, or higher stiffness (parylene-C) can lead to a reduction in the receive performance of PMUTs. We have also shown the impact of the PCB material through simulations and experiments. Although not negatively impacting the overall performance, its presence introduced thickness mode resonance peaks which, at specific frequencies, decreased the receive sensitivity. This result, could serve as a basis for guiding the selection of frequencies to be used, depending on the application. Furthermore, we have also shown that modifications in the measurement setup, could lead to significant measurement errors. In conclusion, this study stands as a foundation for PMUT encapsulation design, presenting a comprehensive analysis of the receive performance of the devices, by evaluating aspects from multiple domains.

## Acknowledgment

We thank Stefan Tornedde, Janine Stockmeyer, and Markus Wöhrmann from Fraunhofer IZM, Berlin, for providing material samples and for the support during MUT assembly, and Okmetic, Vantaa, Finland, for supplying the C-SOI wafers.

